# Synthetic And Natural Face Identity Processing Share Common Mechanisms

**DOI:** 10.1101/2024.08.03.605972

**Authors:** Kim Uittenhove, Hatef Otroshi Shahreza, Sébastien Marcel, Meike Ramon

## Abstract

Recent developments in generative AI offer the means to create synthetic identities, or deepfakes, at scale. As deepfake faces and voices become indistinguishable from real ones, they are considered as promising alternatives for research and development to enhance fairness and protect humans’ rights to privacy. Notwithstanding these efforts and intentions, a basic question remains unanswered: Are natural faces and facial deepfakes perceived and remembered in the same way? Using images created via professional photography on the one hand, and a state-of-the-art generative model on the other, we investigated the most studied process of face cognition: perceptual matching and discrimination of facial identity. Our results demonstrate that identity discrimination of natural *and* synthetic faces is governed by the same underlying perceptual mechanisms: objective stimulus similarity and observers’ ability level. These findings provide empirical support both for the societal risks associated with deepfakes, while also underscoring the utility of synthetic identities for research and development.

## INTRODUCTION

Traditionally, vision research involved the use of analogue images to study level low-, mid-, or high-level visual processes [1, 2, 3]. Over time, novel tools to create, manipulate, and display digital stimuli emerged, which were readily adopted, particularly in the field of face processing. In the early 2000s, commercial photo editing software provided means to achieve even unnoticeable facial manipulations [4]. Early generative software enabled creation of images, which could be readily detected as artificial, i.e. discerned from natural images of real people [5, 6].

### Synthetic Faces: Status Quo

Recent advances in generative AI have fundamentally changed this situation, as well as society at large. In the visual domain, Generative Adversarial Networks (GANs) and Diffusion Models (DMs) enable rapid creation of synthetic facial identities, or deepfakes, at scale [7, 8]. Most importantly, however, the *quality* of deepfakes has increased dramatically. Contrary to early approaches, state-of-the-art (SOTA) models can generate extremely realistic, i.e. natural-looking synthetic identities, and for a given synthetic identity produce images involvinf viewpoint and age-related changes [9].

Mirroring advances in deepfake generation, a growing body of research is focused on machine-based solutions for *automatic deepfake detection* [10]. *Human* deepfake processing on the other hand remains understudied. The limited number of empirical investigations collectively demonstrate that humans cannot reliably distinguish natural from synthetic faces [8, 11, 7, 12]. Law enforcement professionals, who could be tasked with authenticating the veracity of digital context, are challenged by deepfakes – even those with superior face processing skills [11]. Moreover, interventions and training do not lead to significant improvements [13, 14]. Our inability to distinguish natural from synthetic identities harbors both substantial negative and positive potential across different areas of society.

### Challenges and Opportunities

On the one hand, deepfakes can serve as means to deceptive and exploitative ends targeting individuals, e.g., victims of revenge porn and fraud [15, 16]. At the societal level, digital proliferation of AI-generated disinformation campaigns can interfere with processes and decrease trust in governments and institutions [17, 18]. International law enforcement agencies regard synthetic information and deepfakes as a major challenge and a threat to society at large [19, 20]. This situation is compounded by the difficulty in prosecuting deepfakes due to inadequate legal frameworks [17]. A positive effect of the ability to create indistinguishable-from-natural content is the support of creative avenues for artists and media professionals [21]. For example, synthetic avatars are commonplace in the gaming industry and various types of deepfakes are used in the film industry [22].

In the scientific community, considerable efforts are currently directed towards the creation of synthetic face datasets [23, 24, 25, 26, 27, 28], which offer at least two main advantages. First, synthetic datasets are devoid of the legal, ethical, and privacy concerns associated with datasets of real identities, which are often crawled from internet without individuals’ consent, and are used to train face recognition models. Note that several of such web-crawled face datasets, such as VGGFace2 [29] and MS-Celeb-1M [30] have been retracted by their owners due to privacy concerns [31]. Second, synthetic datasets could also save resources, e.g., those associated with the creation of shareable, fair natural databases. Recent advances in SOTA generative models, e.g. [32, 33, 34], can provide tools to generate complex synthetic datasets with sufficient real-life variations. For instance, generative models based on Neural Radiance Fields (NeRF) [35], such as EG3D [34], are capable of 3D identity representations. This allows to generate not only distinct facial identities, but also create any desired viewpoint of a given synthetic identity.

In practice, however, creating synthetic datasets that mirror the variability of natural faces remains a major challenge. Put simply, natural and synthetic faces may differ in terms of their respective feature dimensions, and in turn their theoretical multidimensional face spaces [36]. Realistically, a formal comparison of the *full* range of human and synthetic variability is, of course, not feasible. However, a thorough and fair comparison of subsets of natural and synthetic faces *can* be achieved via in-depth investigation of human performance, under careful consideration of its known determinants.

### Determinants of Natural Face Discrimination

Arguably, the most challenging computations performed by the human brain are those required to discriminate unfamiliar faces [37]. Deciding whether two images show the same, or two different people is an evolutionarily recent and extremely error-prone task [38, 39] —even for highly trained professionals [40]. Observers’ face identity matching performance is determined on the one hand by objective external factors, e.g. stimulus similarity and viewing conditions such as exposure duration and resolution [41, 42, 43]. Moreover, face matching performance will vary between individual observers, due to stable, inter-individual differences in face identity processing (FIP; [44, 45, 46, 47, 48]). Overall, the same external conditions affect individuals differently, depending on their respective location on the FIP ability spectrum [49].

### Synthetic and Natural Face Discrimination: Shared Cognitive Mechanisms?

In this study we sought to answer the currently open question of whether synthetic identity processing is governed by the same mechanisms that determine natural face processing. To this end, we investigated the impact of stimulus similarity and differences in observers’ FIP ability in two experiments probing rapid natural and synthetic face discrimination. Uniquely, we use two different types of face stimuli, derived from best-case real-world and generative scenarios. First, natural identities were represented via professional photographs derived from an artistic project that captured thousands of identities under strictly controlled and thus comparable conditions. Second, synthetic identities were created using a SOTA generative model that can achieve 3D viewpoint-invariant identity representations [34].

We provide the first empirical evidence that FIP for stimuli created with SOTA generative models is governed by the same mechanisms as those involved in natural face discrimination. The results underline the challenges of state-of-the-art facial deepfakes and support the use of synthetic face likenesses as an ecologically valid tool for the study of face processing in general. Filling this knowledge gap contributes to the development of the lacking unified framework of natural information processing accounting for commonalities and differences between humans and machines [37, 50].

## METHODS

All procedures and protocols were approved by the Ethics Committee of the University of Fribourg (approval number 473), the University of Lausanne, and conducted in accordance with the guidelines set forth in the Declaration of Helsinki. All volunteering participants provided informed consent, and were not financially compensated for their participation.

### Participants

Data collection was distributed via students enrolled in a seminar at the University of Lausanne, who recruited up to five participants each. Participants (N=104) completed two face discrimination experiments, in which natural or synthetic stimuli were used, respectively. The majority (n=97) then completed an independent assessment of face identity processing ability (see below). Of these 97 complete data sets, seven were excluded due to exceedingly elevated reaction times across both face discrimination experiments (> 3rd quartile + 1.5* interquartile range, n = 5), or low accuracy (<1st quartile - 1.5* interquartile range, n=2). The final sample subject to analyses comprised 90 participants (41 and 59 identifying as fe-/male, mean age: 32, SD=14, range: 18 to 62), 80 of which were right-handed (7 left-handed, 3 ambidextrous).

### Synthetic and Natural Face Discrimination

All participants completed two face discrimination experiments, which involved the same design, but presented either natural or synthetic face images as stimulus material. These experiments were delivered via Testable [51]; order of completion was counterbalanced across participants. Participants performed 2-alternative forced-choice (2AFC) identity matching for pairs of stimuli presented in rapid succession (see Figure 1), indicating their responses via (bi-manual) button-press. Target and probe stimuli were presented for 500ms each, with a 500ms inter-stimulus interval; observers had unlimited time to time to indicate their decision. Targets’ and probes’ location was centrally offset to prevent the use of local matching strategy (-/+25 pixels vertically and horizontally from the center in four quasi-random locations). Each experiment comprised 420 trials (210 *same/diJerent*), across which different combinations of 21 Caucasian female identities were presented. *Same* trials were generated by pairing five different images of the same identity with one another. *DiJerent* trials were generated by pairing each of the 21 identities with every other identity. Each face discrimination experiment began with four practice trials, which displayed identities not included in the experiment, and which were excluded from analyses.

**Figure 1.**
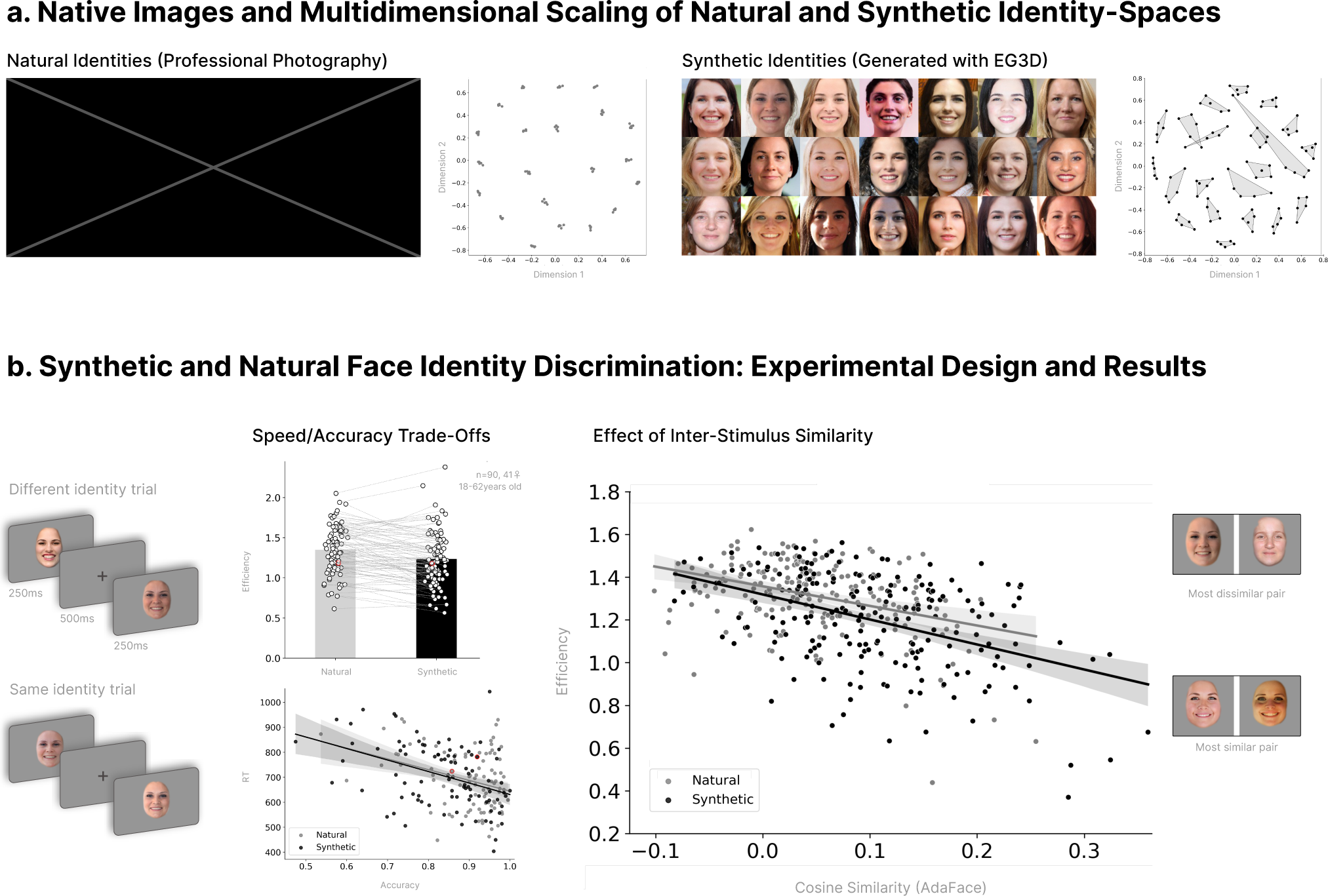
Natural and Synthetic Face Discrimination. **a**. Examples of native stimuli and multi-dimensional scaling of face space. Images of identities were either created via professional photography (left), or generated using EG3D [34], a model based on Generative Adversarial Network and Neural Radiance Fields structure (right).**b**. Schematic of face discrimination experiments and observed relationship between performance measures across experiments. Examples of different and same identity trials (left), and individual observers’ overall efficiency and RT as a function of accuracy across experiments. Effect of inter-stimulus similarity computed using AdaFace on face discrimination efficiency (right) with most dis-/similar image pairs.

#### Natural Face Stimuli

Natural identities were selected from the FACITY image database created and kindly provided by the professional photographer H. Caspar. This database was created as an artistic project, through which hundreds of professional photographers worldwide took images of thousands of peoples’ likenesses following the same criteria (i.e. under the same conditions, to ensure comparable image quality and style; see Figure 1a). From this large database, 21 target identities were selected, all of which were devoid of obvious features (e.g., piercings, moles, scars, etc.) that could aid rapid matching based on specific image features, as opposed to identity matching. Figure 1a shows the set of natural identities used in the natural face discrimination experiment. The selected native images were then processed to create the final stimuli for the natural face discrimination experiment. The midpoint of each face was vertically offset from the center by approximately 10% of the image height. To ensure that natural and synthetic stimuli occupied the same image space in both dimensions, we applied padding to the top of the image (20% of the image height) and to either side of the image (10% of the image width). The resulting images were centered on the midpoint of the face, which (consistent with synthetic images) occupied ca. 55% of the horizontal space. These images were resized to their original size (512×512 pixels, 96 dpi). For each identity we created four modified versions by in-/decreasing the contrast and luminosity of the original image. To prevent the use of external features and contour information, faces were cropped and edges were blurred using the Pyfacer library, with pre-trained models for face detection ([52]) and face parsing ([53]). We created a binary mask to isolate the facial region (removing a section of the top to resolve artifacts around the top of the natural images, due to the nature of their creation in the artistic database). To extract the edges of the facial region mask, we checked each pixel of the mask (value = 1) and marked this pixel as part of the edge if any of the eight surrounding pixels was not part of the facial mask (value = 0). We defined an edge zone by dilating the extracted edge using a 15×15 kernel. For a natural blending effect, we first applied a 29×29 Gaussian kernel to the face region, creating a blurred version. We then recombined the blurred image with the original image: pixels within the edge zone were replaced with pixels from the blurred version, while pixels outside this zone retained their original values. Finally, we introduced further variability by scaling down the face size and applying a rotation to the four additional instances of the original image. Stimuli were presented on a full-screen grey background; a calibration prior to each experiment ensured comparable on-screen stimulus size across devices.

#### Synthetic Face Stimuli

Synthetic identities were generated using EG3D [34]. EG3D is a face generator model based on Generative Adversarial Network (GAN) and Neural Radiance Fields (NeRF) structure. It starts from a random ***z*** *œ N* (0*, I*) to generate an intermediate latent code ***w*** *œ W* that is fed a long with camera parameters to generate a face image from desired point of view. Starting from different random noise, we generated several face identities with frontal pose, retaining only those whose likeness appearance was female Caucasian within the range of ca. 20-40 years. Per synthetic identity, multiple instances were generated by adding random noise to the intermediate latent code ***w*** of the reference instance. The resulting different images per identity could vary in their visual characteristics and facial expressions. Identity correspondence across these images was ensured using the pre-trained face recognition model ArcFace [54] face recognition model. To this end, similar to the typical operation of face verification systems [55, 56], we extracted ArcFace feature vectors from each generated instance and the reference instance. Then, if the cosine similarity of the feature vectors for the generated and reference instance was greater than a predefined threshold, we considered the new instance as the same identity. Only images devoid of paraphernalia (e.g., glasses or hats) and artifacts (e.g, on the skin or eyes) were selected, and processed (cropped, contour-blurred) in the same manner as described above for the natural face images.

#### Multi-Dimensional Scaling of Face Stimuli

The calculation of pairwise similarity between *n* stimuli places each stimulus in an *n*-dimensional space where its position is determined by its dis-/similarity to all other stimuli. Multi-dimensional scaling (MDS) allows us to create a lower-dimensional embedding to visualize the dis-/similarity between stimuli in a simple way, as shown in Figure 1a. First, we transformed our similarity matrix into a dissimilarity matrix by subtracting the similarity values from 1. Then, we created a two-dimensional embedding using the MDS function from the scikit learn manifold package [57]. Larger distance between points reflects more dissimilarity between the stimuli. Stimuli representing the same identity are closer in space (especially for natural stimuli; see above); for synthetic stimuli, convex hulls encapsulate points indicate stimuli representing the same identity.

### Assessment of Participants’ Face Identity Processing Ability

Participants were invited to participate in independent tests of face perception and memory ability. The tests included the long version of the Yearbook Test (YBT; [58, 45]), and the Cambridge Face Memory Test (CFMT+; [59]). The YBT probes face perception through unfamiliar face identity matching across image variations and considerable age-related changes in appearance. Participants are presented with 8 (same-gender) panels displaying five (young adult) target identities and 10 (older) probes, half of which represent non-match distractor identities. The CFMT+ assesses face memory by soliciting identity recognition for six experimentally learned male Caucasian identities in a 3-alternative forced-choice format under increasingly difficult visual conditions. These tests represent sensitive means to assess individual differences in observers’ face identity processing ability, and are used to identify so-called Super-Recognizers, i.e. individuals with naturally superior ability for processing facial identity [46, 11, 60, 61]. The YBT was administered as a paper-based test (cf. [58, 45], while the CFMT+ was administered online via Testable [51].

### Analyses

Our analyses aimed to answer the question of whether perceptual discrimination of, i.e. telling apart synthetic identities follows the same pattern as that of natural ones. In a first step, we omitted trials associated with exceedingly short (<250ms) or long (> third quartile + 1.5* inter-quartile range) RTs (considered to reflect anticipatory responses or external disruption). This led to elimination of an average of 6.81% of RTs/trials per participant (SD=2.43%, range=2.02-18.93%). To investigate face *discrimination* in the natural (*FD_nat_*) and synthetic (*FD_syn_*) experiments, we analyzed *diJerent* trails to determine the relationship between performance measures (accuracy, RTs, efficiency) and objective inter-stimulus similarity, and individual observers’ independently assessed FIP ability, respectively. Bayesian analyses were performed with JASP version 0.18.3, following the approach recommended by [62] that allows quantifying evidence for the null and alternative hypotheses. Bayes Factors (BFs) were interpreted as follows: values between 1/3 and 3 were considered inconclusive, values from 1/10 to 1/3, or 3 to 10 indicated substantial evidence, and values less than 1/10 or greater than 10 indicated strong evidence. To maintain neutrality, we used non-informative priors in all analyses, allowing the data to drive our inferences. Specifically, we conducted Bayesian linear regressions using a JZS prior with an r scale of .354 for the coefficients and a Beta Binomial model prior with parameters a = 1 and b = 1. Our aim was to quantify the evidence for the inclusion of each predictor across different models. For comparisons between conditions, we used a Bayesian Wilcoxon signed-rank test with a Cauchy prior with scale of .707.

#### Overall comparison of performance for Natural and Synthetic Face Matching: Efficiency and Speed/Accuracy Relationships

First, we compared individual observers’ efficiency scores (accuracy/RT*1000) across experiments. We investigated the nature of the *relationship* between accuracy and speed across experiments and participants using a Bayesian linear regression on the log of RT as dependent variable, and with accuracy and experiment (natural vs. synthetic), as well as their interaction, as predictors. Here, a similar relationship is taken to suggest that synthetic and natural faces were generally processed in a comparable manner.

#### Objective Stimulus Similarity and Observers’ Independently Assessed Face Identity Processing Ability

In a second step, we investigated the relationship between observers’ performance for the *FD_nat_* and *FD_syn_* experiments in greater detail. We examined observers’ performance measures as a function of the pairwise similarity between identities as determined using the pre-trained face recognition model AdaFace [63]. AdaFace computes the cosine similarity between the features extracted from the images of each trial (i.e, image pair), resulting in a similarity score ranging from -1 (maximum dissimilarity) to 1 (maximum similarity). This procedure has been used in face verification systems to match a pair of face images. We created similarity score bins of 0.05 and calculated observers’ efficiency per bin. Additionally, we determined whether *FD_nat_* and *FD_syn_* identity discrimination performance systematically varied depending on observers’ overall face identity processing (FIP) ability. This overall ability was computed based on their performance across two sensitive tests of face *matching* and *recognition*. To this end we computed each observer’s overall FIP ability scores by averaging their individual z-standardized YBT and CFMT+ scores.

## RESULTS

### Natural and Synthetic Face Discrimination: Overall Performance

Bayesian Wilcoxon signed-rank test of individual observers’ aggregate scores of efficiency yielded strong evidence in favor of an effect of *experiment* (BF10=77, W=3166, Rhat=1.017). Bayesian linear regression with log *RT* as dependent variable and *accuracy* as independent variable, with *experiment* as moderator, indicated strong evidence for an effect of *accuracy* on *RT* (coefficient: -.256; 95% CI: -.351, -.139; BFinclusion=11624), reflecting higher accuracy being associated with lower RTs. The analysis yielded evidence against an effect a main effect of *experiment* (coefficient: .0003; 95% CI: -.072, .036; BFinclusion=.278), and against a moderating effect of *experiment* (coefficient: -.002; 95% CI: -.117, .006; BFinclusion=.233). Overall, we find evidence against (differential) speed/accuracy trade-offs across experiments.

### Effects of Objective Similarity and Observers’ Face Processing Ability

Figure 1b displays overall performance for trials showing different identities for each *experiment*, and as a function of stimulus *similarity*. Bayesian linear regression analysis of efficiency scores indicated evidence for an effect of *experiment* (coefficient: -.045; 95% CI: [-.089, .000]; BFinclusion=3.270), stimulus *similarity* (coefficient: -.831; 95% CI: [-1.017, -.673]; BFinclusion=6.788 *·* 10^23^), and FIP *ability* (coefficient: .040; 95% CI: [*≠*5.164 *·* 10^6^, .067]; BFinclusion=44.113). We found inconclusive evidence against an interaction between FIP and experiment (*ability* x *experiment*, BFinclusion=.383). Critically, we found evidence against (any) interactions involving the factor *similiarity* (*similarity* x *experiment*, BFinclusion=.165; *ability* x *similarity*, BFinclusion=.208; *ability* x *experiment* x *similarity*, BFinclusion=.034).

## DISCUSSION

We investigated whether natural and synthetic facial identity discrimination is governed by the same perceptual mechanisms. Uniquely, we used images derived from large-scale databases comprising thousands of identities derived from either professional photography of real people, and an open source SOTA generative model optimized for viewpoint invariance [34]. For both natural and synthetic images, we quantified the similarity between identities and image pairs [63], and assessed observers’ individual FIP abilities.

To our knowledge, the present study provides the first empirical evidence that FIP observed for natural facial identities extends to stimuli created with SOTA generative models. Filling this knowledge gap provides three crucial contributions. First, it contributes to the development of the lacking unified framework of natural information processing accounting for commonalities and differences between humans and machines [37, 50]. Second, it underscores the major societal challenges associated with SOTA facial deepfakes and their detection [8, 11]. Third, and finally, it supports the use of synthetic face likenesses as an ecologically valid tool for research on face processing and its applications.

### Comparable Processing of Natural and Synthetic identities

Across experiments, we report strong evidence that that both objective stimulus similarity, as well as individual differences in observers’ FIP abilities affect natural and synthetic face discrimination in the same manner. Observers’ face discrimination performance varied with their independently measured FIP ability, but their performance did not vary as a function of face type. Confronted with a pair of different identities – natural or synthetic – observers’ exhibited comparable face discrimination efficiency.

Moreover, across face types and observers, we found comparable effects of objective stimulus similarity. Performance decreased with increased inter-identity similarity, and observers with higher FIP ability showed higher proficiency. To our knowledge, these results provide the first, and moreover compelling evidence that synthetic faces are indeed processed in the same manner as natural ones. These findings emerged in the context of stimuli created using a generative model that enables viewpoint variations and dynamic content of a given synthetic facial identity [34]. These features provide a range of advantages across research and applied settings, e.g. comparative assessment of human and machine performance under challenging, simulated surveillance footage conditions.

### Outlook

Our findings suggest that synthetic identities can, in principle, be generated and used as alternatives where, traditionally, natural faces were used. In research settings, synthetic face stimuli could entirely replace the use of images of human likenesses. The advantages are clear: The absence of privacy concerns enable sharing and thus generally contribute to enhancing reproducibility and replicability of research. Generative models offer the means to enhance fairness, e.g. by creating synthetic datasets that are balanced in terms of demographic diversity.

The advantage of synthetic faces extends to applied areas, particularly where comparable assessment of individuals is critical. Shareable synthetic datasets enable comparable testing of abilities for high stakes professional roles, e.g. forensic expert training and continued performance evaluation. Law enforcement professionals dealing with eye witness testimonies could use synthetic image lineups, as opposed to images of individuals who e.g. had been taken into custody previously. Rather than having to locate a specific identity with certainty, witnesses could select identities they perceived as most similar to the perpetrator. The selected probable synthetic matches could then be compared to criminal databases. This would offer enhanced protection of former criminals’ privacy (e.g., to avoid potential stigmatization), as their identity would no longer be revealed to unrelated victims.

Notwithstanding their advantages, synthetic faces are not a simple, immediately ready-to-use alternative. After automated model-based stimulus generative, manual selection of appropriate images is always required. In our case, we generated numerous iterations of identity pools, from which target identities were selected. Indeed, a very large proportion of synthetic identities had to be excluded due to the presence of abnormal information (see Supplemental Material for examples of exclusions). Further work is required to determine whether the similarities we report for frontal natural and synthetic face matching — akin to mugshot comparisons — extends to other real-life scenarios. For instance, are model-based variations of synthetic identities (e.g., ageing, viewpoint, illumination) perceived in the same way as for natural faces? Our study also provides a canvas to determine whether processing of natural/synthetic identity *voices* also involves shared mechanisms.

## CONCLUSION

Synthetic identities are ubiquitous and will remain a fixture of digital societies. We believe that synthetic identities provide a useful tool for future research, and across a broad range of applications. They offer means to increase transparency and privacy protection alike. Outwith academia and industry, their advantages should be leveraged in the design of equitably accessible AI literacy programs, culturally tailored for its recipients.

## ACKNOWLEDGEMENTS

MR is grateful to H. Caspar for kindly sharing his art. KU and MR thank their previous students for support during data acquisition and all volunteering observers for their participation. MR is supported by a Swiss National Science Foundation PRIMA (Promoting Women in Academia) grant (PR00P1 179872). HOS is supported by the H2020 TReSPAsS-ETN Marie Sk≥odowska-Curie early training network (grant agreement 860813); SM is supported by the Idiap Research Institute.

## AUTHOR CONTRIBUTION STATEMENT

MR: conceived and planned the experiments; wrote all sections of the manuscript; supervised and interpreted findings; created visualizations. KU: prepared stimuli and experiment; supervised data acquisition; managed data; wrote parts of the methods and results section; conducted statistical analyses, created visualizations; created and supplied the OFS repository. HOS and SM provided the synthetic face stimuli, pre-processed all stimuli used in the natural and synthetic face discrimination experiments, extracted the similarity scores, wrote parts of the methods and contributed to discussions. All authors approved the final version of the manuscript.

## DATA, MATERIAL, AND SOFTWARE AVAILABILITY

Anonymized research data reported subject to analysis and analysis code can be found on the accompanying OSF project.

## SUPPLEMENTARY MATERIAL

### Examples of Discarded Generated Synthetic Identities

We initially generated more than a thousand synthetic identities and selected 128 identities which looked Caucasian female with middle age. Then, by resampling around the corresponding latent vector, we generated different instances for each identity from a frontal viewpoint. However, many of these identities and their associated instances contained paraphernalia and artifacts, which made them unsuitable for our study. The high prevalence of artifacts may result from the rendering process, whereby a lower-resolution 3D embedding was decoded into higher-resolution 2D viewpoints, which may have introduced inconsistencies or unnatural textures in some cases. Ultimately, only 30 identities and their corresponding instances met our criteria for suitability. The figure below illustrates examples of the discarded identities.

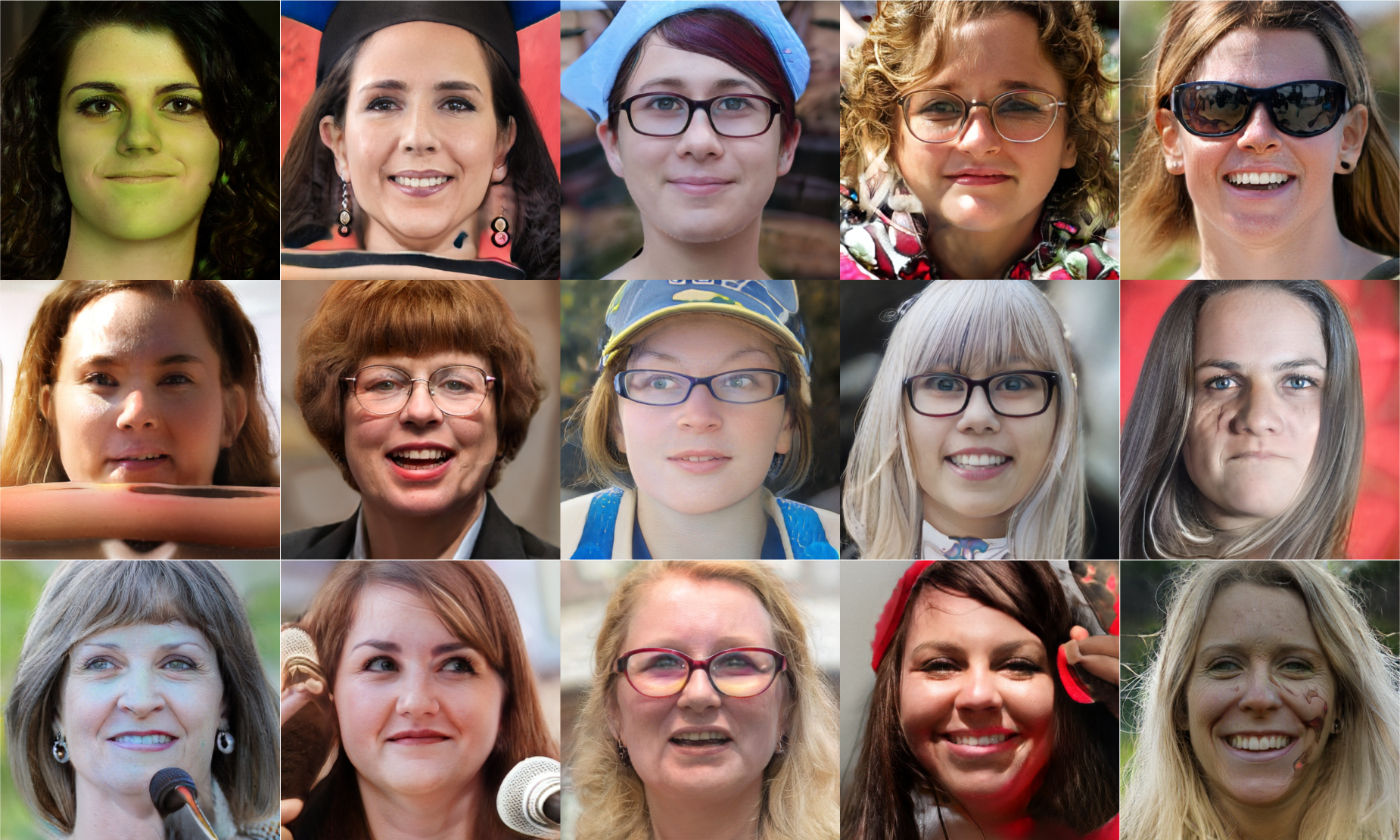

## Notes

### Competing Interest Statement

The authors have declared no competing interest.

